# Quiescent cells actively replenish CENP-A nucleosomes to maintain centromere identity and proliferative potential

**DOI:** 10.1101/433391

**Authors:** S. Zachary Swartz, Liliana S. McKay, Kuan-Chung Su, Abbas Padeganeh, Paul S. Maddox, Kristin A. Knouse, Iain M. Cheeseman

## Abstract

Centromeres provide a robust model for epigenetic inheritance as they are specified by sequence-independent mechanisms involving the histone H3-variant CENP-A. Prevailing models indicate that the high intrinsic stability of CENP-A nucleosomes maintains centromere identity indefinitely. Here, we demonstrate that CENP-A is not stable at centromeres, but is instead gradually and continuously incorporated in quiescent cells including G0-arrested tissue culture cells and prophase I-arrested oocytes. Quiescent CENP-A incorporation involves the canonical CENP-A deposition machinery, but displays distinct requirements from cell cycle-dependent deposition. We demonstrate that Plk1 is required specifically for G1 CENP-A deposition, whereas transcription promotes CENP-A incorporation in quiescent oocytes. Preventing CENP-A deposition during quiescence results in significantly reduced CENP-A levels and perturbs chromosome segregation following the resumption of cell division. In contrast to quiescent cells, terminally differentiated cells fail to maintain CENP-A levels. Our work reveals that quiescent cells actively maintain centromere identity providing an indicator of proliferative potential.

## Introduction

Heritable information is propagated by DNA sequence as well as via sequence-independent epigenetic marks that control the properties or activity of specific genomic loci. These marks include covalent changes to the DNA itself, such as methylation, post-translational modifications to histone proteins, and the incorporation of histone variants such as H2A.Z, macroH2A, and the histone H3 variant CENP-A. For an epigenetic mark to stably alter the behavior of a locus, it must be restricted to the correct location, propagated to new cells formed during cell division, and maintained under all circumstances where this information is needed to direct cellular behaviors. To understand the basis for epigenetic specification, it is critical to determine how these marks are stably maintained for extended periods. We sought to understand these requirements by focusing on the epigenetic specification that occurs at centromeres.

Inheritance of genetic information across cell divisions (to every cell in the body) and generations (from parent to progeny) requires the presence of the centromere at a single locus on each chromosome (McKinley and Cheeseman, 2016). Centromeres serve as the foundation for assembly of the macromolecular kinetochore structure that connects chromosomes to the spindle apparatus to direct their segregation (Cheeseman, 2014). Specific DNA sequences are neither necessary nor sufficient for centromere identity and function in most organisms, and instead centromeres are defined epigenetically by the presence of the histone H3-variant CENP-A. CENP-A is required for the localization of all known kinetochore components such that the loss of CENP-A results in a failure of kinetochore function, chromosome mis-segregation, and ultimately cell inviability or dysfunction. Thus, the central requirement for centromere identity and function is to specify, propagate, and maintain the presence of CENP-A nucleosomes at a single site on each chromosome.

Pre-existing CENP-A nucleosomes are required to direct new CENP-A deposition (McKinley and Cheeseman, 2016) and de novo centromere formation is exceptionally rare (Shang et al., 2013). As such, it is essential that this CENP-A mark be stably retained under all circumstances where subsequent division is required to ensure proper genome inheritance. Current models suggest that CENP-A is indefinitely stable at centromeres, with the replenishment of CENP-A nucleosomes following DNA replication restricted to the following G1 phase (Jansen et al., 2007). Prior work has proposed that pre-existing CENP-A nucleosomes are maintained through their immobility and stability conferred by its binding partners (Bodor et al., 2013; Cao et al., 2018; Guo et al., 2017; Jansen et al., 2007; McKinley and Cheeseman, 2016; Smoak et al., 2016). However, most work has focused on the mechanisms that propagate CENP-A in rapidly dividing mitotic cells. Much less is known about CENP-A maintenance at centromeres in the diverse metazoan cell types that exit the cell cycle for extended periods.

Given the critical role of quiescent cells in organism development and organ homeostasis and repair, it is critical to understand how centromere identity is maintained under diverse physiological situations, including during prolonged cell cycle arrest. Here, we explore CENP-A nucleosome dynamics in quiescent germline and somatic cells across animal species. In contrast to previous reports, we find that CENP-A levels are not indefinitely stable. Our work suggests that pre-existing CENP-A nucleosomes are destabilized in a Pol II-dependent manner and, in quiescent cells, CENP-A is gradually re-incorporated using its canonical deposition machinery to maintain steady state levels. We find that ongoing CENP-A deposition is essential to maintain centromere function and maintain proliferative capacity during quiescence. In contrast, terminally differentiated muscle cells fail to maintain CENP-A levels. This work defines a critical mechanism by which an essential epigenetic mark is maintained in quiescent cells to retain proliferative potential.

## Results

### CENP-A exchanges at centromeres in quiescent human cells

Prior work has analyzed CENP-A deposition in rapidly dividing cells, including human tissue culture cells. In contrast, little is known regarding CENP-A dynamics in non-dividing cells. Therefore, we first asked whether the levels of endogenous CENP-A remain constant in human hTERT RPE-1 tissue culture cells induced to enter G0 by serum starvation and contact inhibition. Based on immunofluorescence against endogenous CENP-A, we found that CENP-A levels at centromeres remained constant across a 14-day time course (Fig. 1A,B). This finding is consistent with either highly stable CENP-A nucleosomes, as suggested by prior work, or with balanced CENP-A loss and incorporation. To distinguish between these possibilities, we introduced a HaloTag homozygously into the endogenous CENP-A locus in RPE-1 cells (Fig. S1A). The HaloTag undergoes covalent conjugation to fluorescent small molecule derivatives, thereby enabling fluorescent pulse/chase-based assays to label and distinguish pre-existing versus newly deposited CENP-A (Fig. 1C). Strikingly, we found that new CENP-A-Halo was gradually incorporated at centromeres in quiescent RPE-1 cells (Fig. 1C,D) despite the absence of division. By normalizing the level of new CENP-A relative to total CENP-A, we found that CENP-A deposition occurred at a rate of approximately 10% of total CENP-A per day, reaching an average of 68% of total CENP-A by day 7 (Fig. 1D). Similar results were obtained when quiescent cells were grown continuously in the presence of the Kif11 inhibitor, STLC, to prevent any rare mitotic divisions (Fig. S1B). Because the total level of CENP-A at centromeres remained unchanged during a 14-day quiescent arrest (Fig. 1A), we conclude that gradual CENP-A exchange occurs at centromeres to maintain constant CENP-A levels in the absence of division.

**Figure 1.**
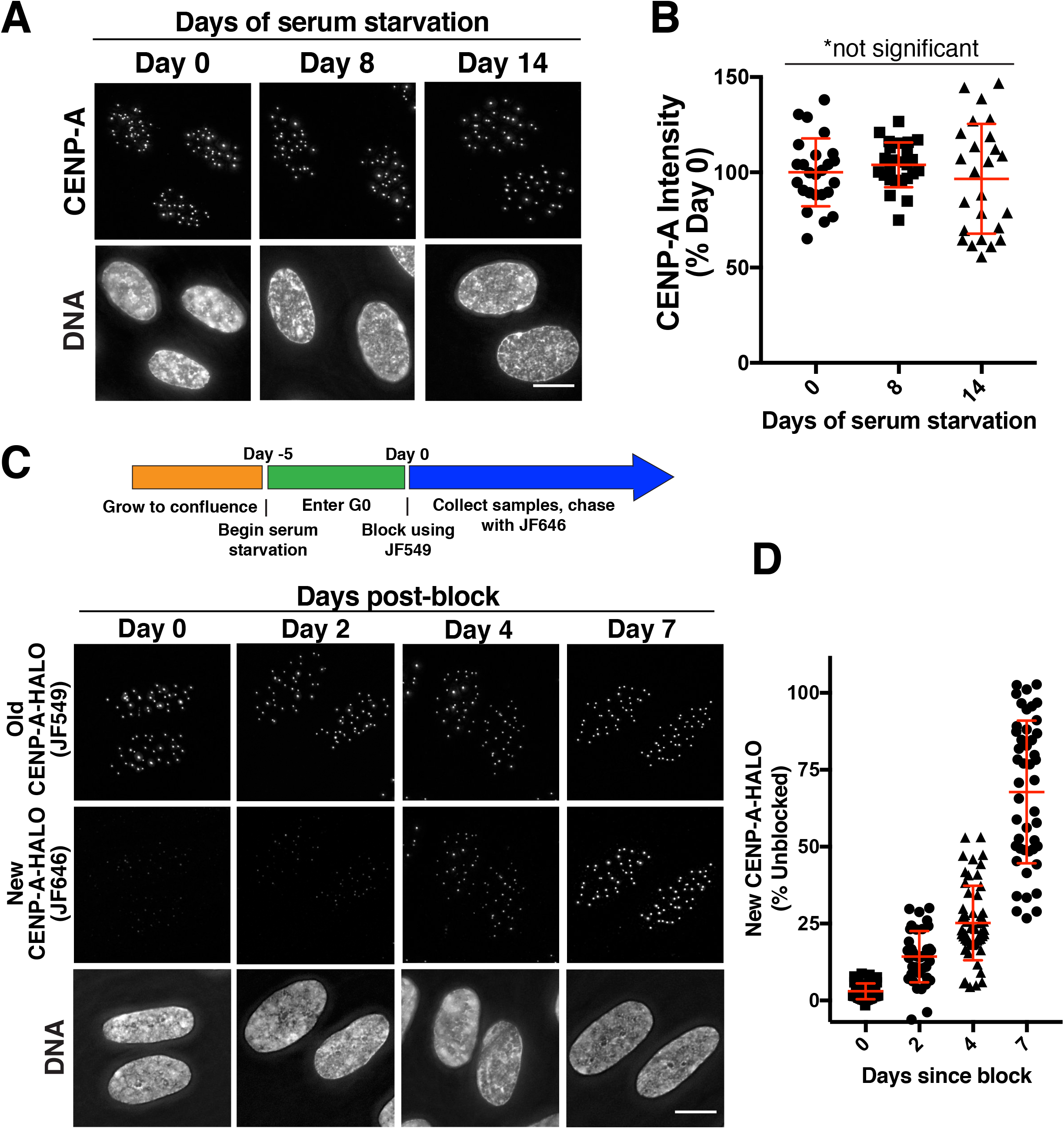
CENP-A is gradually incorporated in quiescent human somatic cells. (A) Immunofluorescence images showing endogenous CENP-A (scaled identically) in quiescent RPE-1 cells at 0, 8, and 14 days after entering serum-starvation induced quiescence. (B) Fluorescence quantification reveals that CENP-A levels remain constant during quiescence. Points represent the sum of all centromeres from an individual cell relative to Day 0. Error bars represent the mean and standard deviation of 25 cells/time point. The differences between time points are not statistically different based on a Welch’s t test. (C) Halo-tag CENP-A incorporation assay in quiescent cells. Schematic indicates experimental conditions. Images - Top: Old HALO-CENP-A visualized with JF549. Middle: New CENP-A-Halo visualized with JF646. Bottom: Corresponding DNA staining. (D) Quantification of new CENP-A-Halo fluorescent intensity relative to pre-existing CENP-A (see Experimental Procedures). Points represent the sum of all centromeres of individual cells. Error bars represent the mean and standard deviation of at least 40 cells. Scale bars = 10 μm.

### CENP-A deposition in quiescent cells requires HJURP and the Mis18 complex

We next sought to define the requirements for deposition of CENP-A at centromeres in quiescent human cells. For these experiments, we used an inducible CRISPR/Cas9-based knockout system in RPE-1 cell lines (McKinley and Cheeseman, 2017; McKinley et al., 2015). Consistent with prior results, the inducible knockout of the CENP-A-specific chaperone HJURP or the Mis18 complex subunit Mis18β resulted in a dramatic reduction in centromeric CENP-A levels and mitotic defects after 7 days of continuous proliferation (Fig. S1C,D). To test the role of HJURP and the Mis18 complex in CENP-A deposition during quiescence, we next induced the knockout as cells entered quiescence (Fig. 2A). To ensure no further divisions occurred, the cells were cultured in the presence of STLC. Uninduced control cells retained CENP-A at centromeres throughout the time course (Fig. 2B,D). In contrast, in HJURP and Mis18β knockout cells, CENP-A centromere levels began declining at day 4 of their quiescent arrest, decreasing by an average of ~30% at day 10 despite the absence of division (Fig. 2B-E). We note that the nature of the inducible Cas9 DNA cleavage results in a mixed population with ~50% of the cells displaying a complete homozygous knockout (see (McKinley and Cheeseman, 2017)), thereby likely under-representing the extent of this effect. To assess the consequences of this reduction in CENP-A levels, we induced cell cycle reentry by serum addition and monitored chromosome segregation in the first mitotic division following release. We found that HJURP inducible knockout cells displayed a significant increase in lagging chromosomes during anaphase, a rigorous measure of chromosome segregation fidelity (Fig. 2F). Thus, the presence of CENP-A nucleosomes at centromeres is maintained by an HJURP and Mis18 complex-dependent mechanism during quiescence in G0-arrested human cells, which is required for proper chromosome segregation following the return to growth.

**Figure 2.**
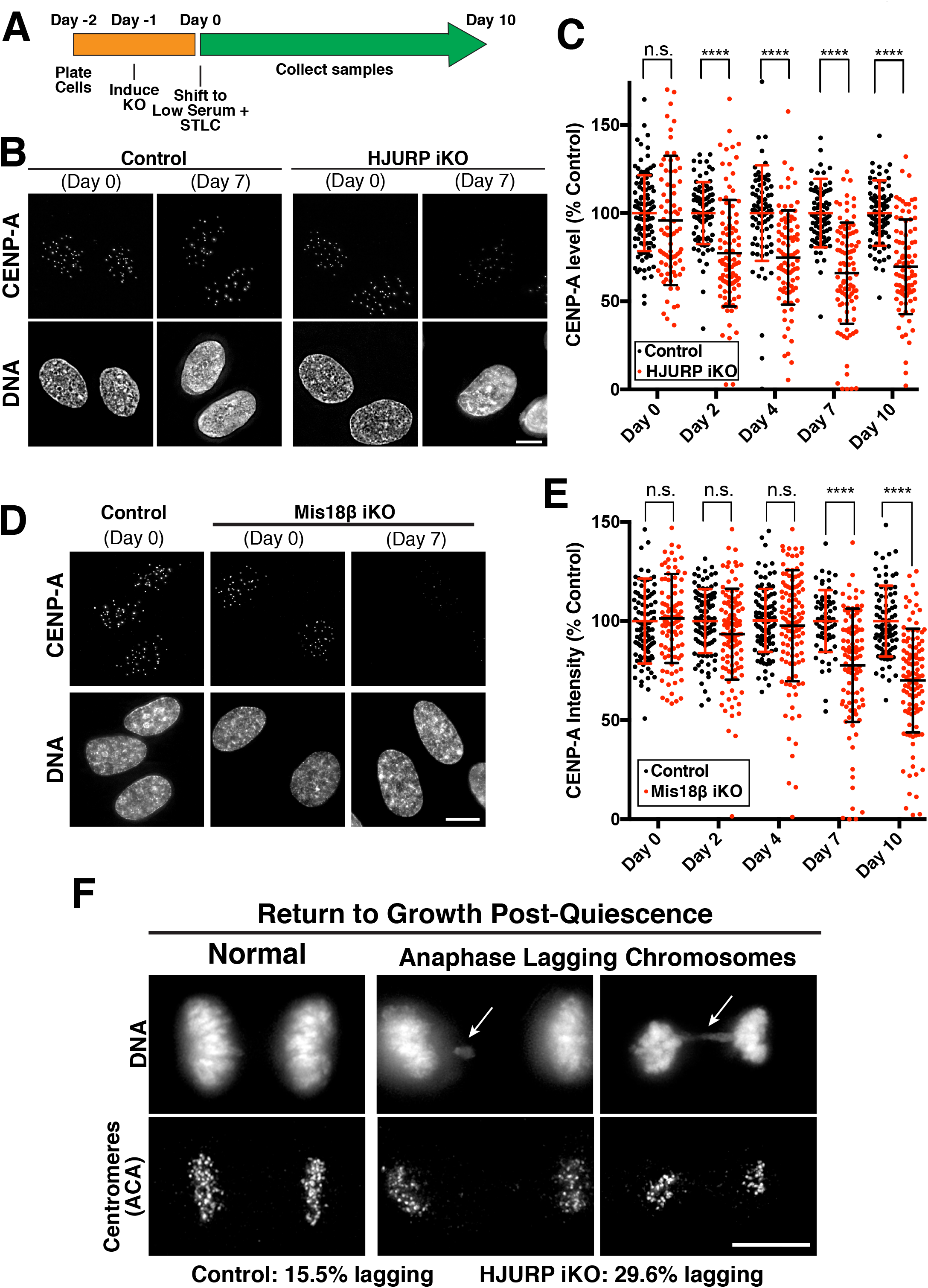
Gradual CENP-A incorporation in quiescent cells requires HJURP and the Mis18 complex. (A) Schematic showing knockout strategy. Knockout of HJURP or Mis18|3 was induced in RPE-1 cells as they entered quiescence, which were then cultured in the presence of the KIF11-inhibitor STLC to ensure no further division. (B) Immunofluorescence for endogenous CENP-A showing a marked reduction in its centromere levels in HJURP knockout cells under quiescent conditions. Scale bar = 5 μm. (C) Fluorescence quantification of centromeric CENP-A in HJURP knockout cells relative to control levels. Each point represents the average of all centromeres within a cell. Error bars represent the mean and standard deviation (Day 0: control n=104, iKO n=81; Day 2: control n=83, iKO n=93; Day 4: control n=80, iKO n=86; Day 7: control n=81, iKO n=93; Day 10: control n=88, iKO n=87 cells). **** p < 0.0001 by two-tailed Mann-Whitney test. (D) Immunofluorescence for CENP-A in the Mis18β knockout, as in (B). (E) Quantification of CENP-A intensity for the Mis18β knockout, as for (C). Scale bar = 5 μm. (Day 0: control n=104, iKO n=95; Day 2: control n=98, iKO n=110; Day 4: control n=106, iKO n=113; Day 7: control n=62, iKO n=101; Day 10: control n=90, iKO n=109 cells). **** p < 0.0001 by two-tailed Mann-Whitney test. (F) Immunofluorescence images showing anaphase chromosome segregation following release from quiescence by serum addition. The presence of lagging chromosomes in anaphase (arrows) was scored as a rigorous measure of chromosome mis-segregation (p = 0.0071 based on a Chi-square test). Control n = 116 anaphase cells; HJURP iKO n = 159 anaphase cells. Scale bar = 10 μm.

### CENP-A is gradually incorporated at centromeres in prophase I-arrested oocytes

The ongoing CENP-A incorporation that we observed in quiescent human tissue culture cells prompted us to test whether this behavior occurs in other cell types that undergo a prolonged cell cycle arrest. Oocytes remain arrested in prophase I of meiosis for extended periods until hormonal stimulus triggers their re-entry into meiosis. As fertilized oocytes divide to create an entire organism, it is critical to maintain centromere identity during this extended prophase I arrest to act as templates for all subsequent centromeres. To define the mechanisms that maintain CENP-A during quiescence, we analyzed oocytes from the starfish *Patiria miniata*, which provide a robust model for cell biological studies of meiosis and early development (Fig. 3A; Borrego-Pinto et al., 2016a; Borrego-Pinto et al., 2016b). To assess centromere dynamics during the meiotic divisions, we identified starfish orthologues of CENP-A and the CENP-A-binding partner, CENP-N. When expressed as GFP fusions by mRNA injection into oocytes, both 3xGFP-CENP-A and 3xGFP-CENP-N localized specifically to centromeres in blastula-stage embryos in which cells undergo frequent divisions (Fig. S2A,B).

**Figure 3.**
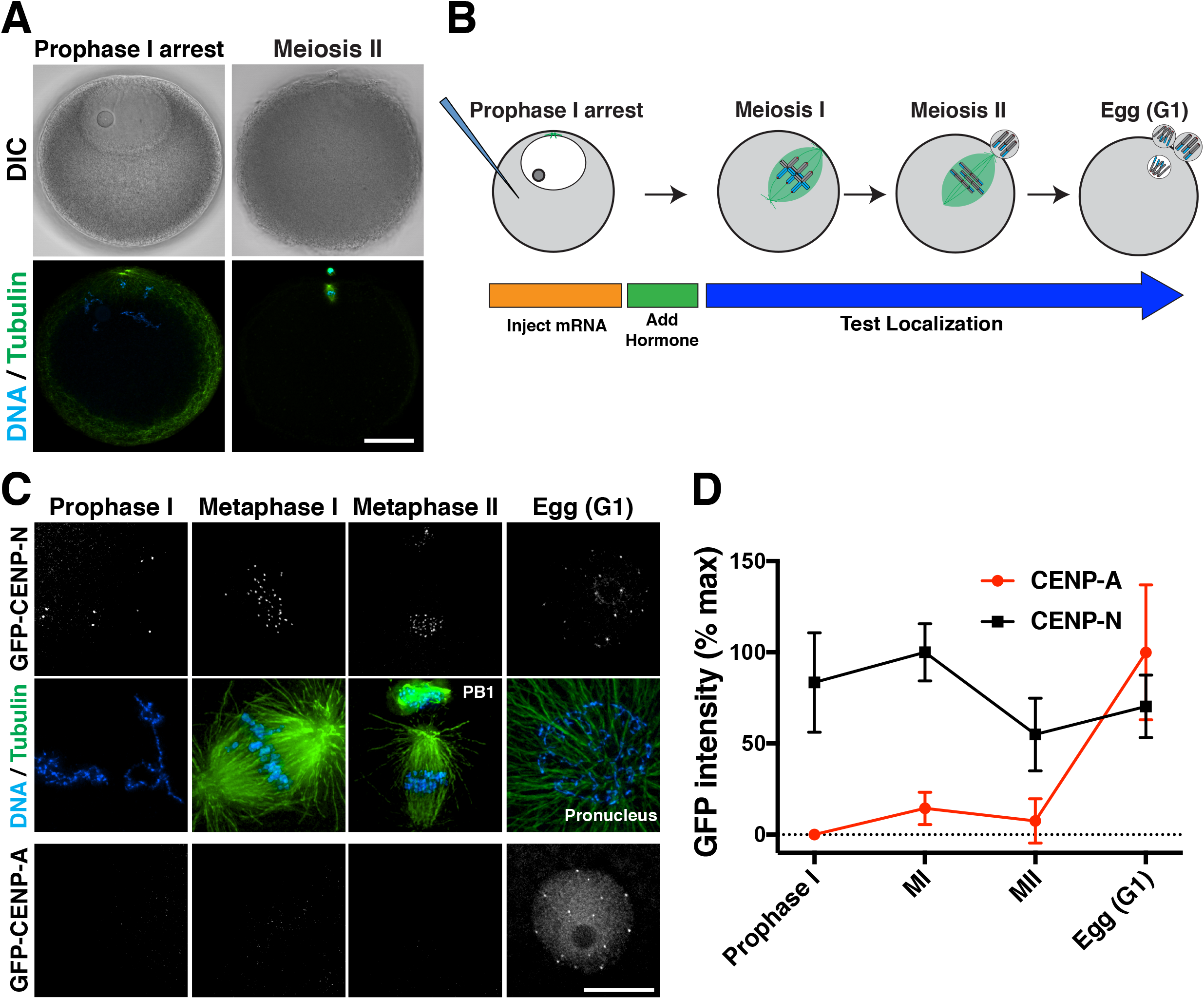
CENP-A is deposited in G1 following meiotic completion. (A) Confocal z-projections of *Patiria miniata* oocytes in prophase I arrest (left) or meiosis II (right), with DIC images (top) and immunofluorescence for microtubules and DNA (bottom). Scale bar = 40 μm. (B) Schematic of meiotic progression and strategy to assess new centromere protein incorporation in starfish oocytes. (C) Localization dynamics for 3xGFP-CENP-N (middle row) and 3xGFP-CENP-A (bottom row). Counter staining for microtubules and DNA is provided on the top row, corresponding to CENP-N oocytes. Microtubules were scaled non-linearly to show cell state. CENP-A and CENP-N images were individually scaled linearly. Scale bar = 10 μm. (D) Quantification of relative CENP-N and CENP-A fluorescent intensity at centromeres throughout meiosis. Each time point represents the mean and standard deviation of at least 6 oocytes.

We next assessed the timing for the localization of newly synthesized centromere proteins in oocytes (Fig. 3B). Starfish oocytes remain arrested in prophase I of meiosis until hormonal stimulation. Upon hormonal stimulation, oocytes undergo two sequential rounds of chromosome segregation (meiosis I and II) without an intervening S-phase. Following mRNA injection of prophase I-arrested oocytes, newly expressed 3xGFP-CENP-N localized to centromeres within 24 hours following mRNA injection (Fig. 3C,D), indicative of rapid turnover. In contrast, new 3xGFP-CENP-A did not localize to centromeres in arrested oocytes (Fig. 3C,D), although soluble protein fluorescence was detected and Western blotting indicated the presence of 3xGFP-CENP-A (Fig. S2C). When meiosis was induced hormonally, we did not observe CENP-A incorporation until completion of meiosis and formation of the female pronucleus (Fig. 3C,D). This timing coincides with the first G1-like cell cycle state following the oocyte-to-embryo transition (Mori et al., 2006; Tachibana et al., 1997). Thus, rapid CENP-A deposition occurs following meiotic exit into G1 in starfish oocytes, similar to the G1 deposition observed in mitotically-dividing human tissue culture cells (Jansen et al., 2007).

As with mammalian oocytes that can remain arrested for decades, starfish oocytes are held in an extended prophase arrest for months until spawning. To assess CENP-A dynamics during the prophase I arrest, we developed a method to culture starfish oocytes in vitro using starfish coelomic fluid, which maintained their competency for meiosis and fertilization for several weeks. Although we did not detect 3xGFP-CENP-A at centromeres at early time points, after 5 days we observed its incorporation at punctate foci (Fig. 4A,B; Fig. S3A). This newly deposited CENP-A was retained at centromeres after meiotic entry and co-localized with antibodies against the centromere protein CENP-C (Fig. 4A; Fig. S3A), indicating that it was incorporated specifically at centromeres. By quantifying 3xGFP-CENP-A levels incorporated during the arrest relative to the level of 3xGFP-CENP-A that is incorporated in the female egg G1 pronucleus, we found that 3xGFP-CENP-A intensity increased gradually at a rate of ~2% per day (Fig. 4B), consistent with a low level of ongoing deposition. New CENP-A incorporation occurred stochastically, with some centromeres in an individual oocyte incorporating >60% of the CENP-A levels deposited during the G1/pronucleus deposition event, but others incorporating relatively little (Fig. S3B). Thus, similar to the behavior of G0-arrested human cells, new CENP-A molecules are gradually incorporated in starfish oocytes during prophase I arrest.

**Figure 4.**
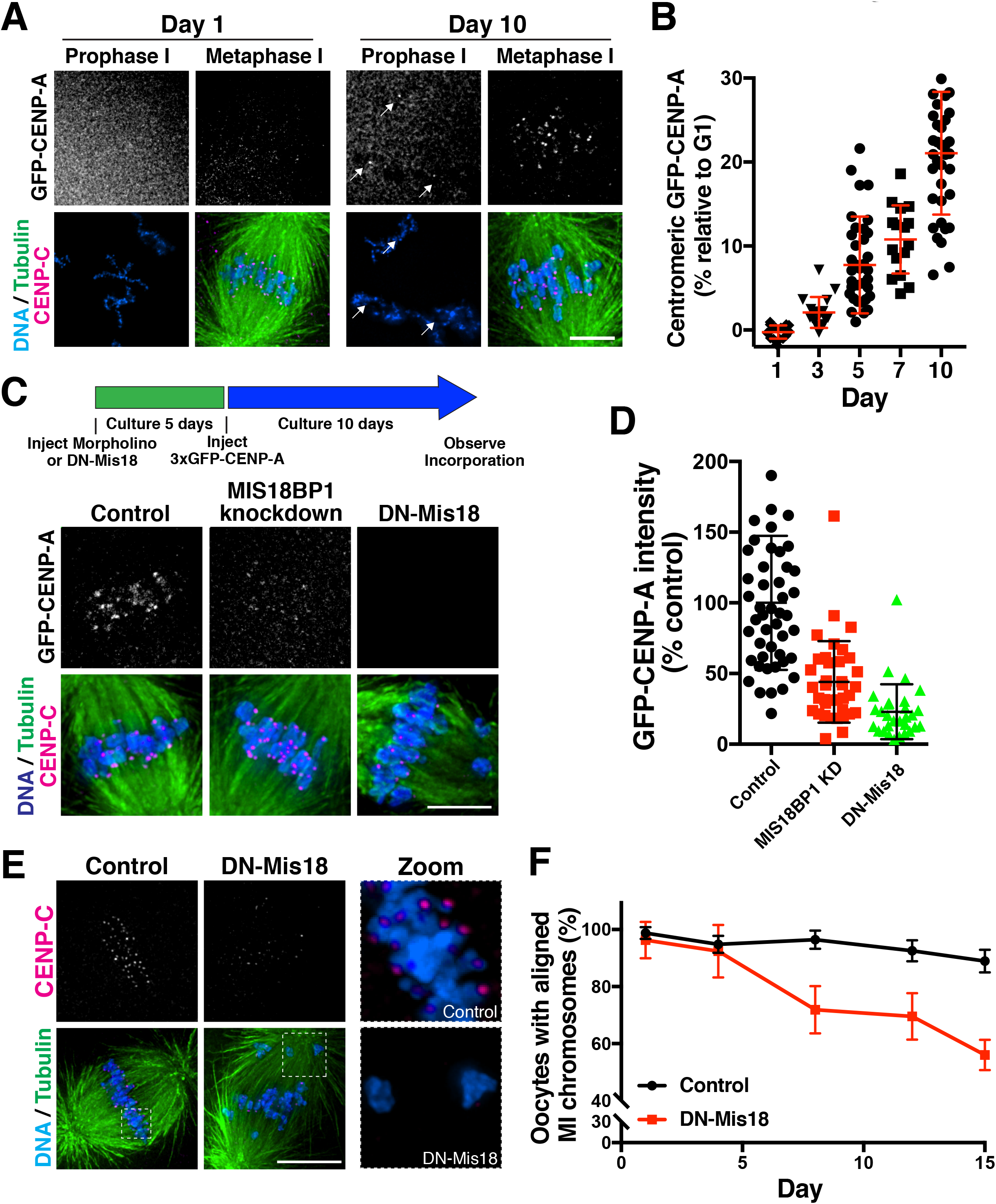
CENP-A is gradually deposited in prophase arrested oocytes. (A) Prophase-arrested chromatin (left) and meiosis I spindles (right) imaged 1 day or 10 days following expression of 3xGFP-CENP-A. 3xGFP-CENP-A images were scaled individually for easier visualization. Microtubule staining was scaled nonlinearly. (B) Quantification of 3xGFP fluorescence intensity incorporated in prophase relative to that deposited in G1 (see Experimental Procedures). Points show the mean of all centromeres from an individual oocyte. Error bars represent the mean and standard deviation of at least 16 oocytes. (C) Prophase incorporation of 3xGFP-CENP-A is reduced following Mis18BP1 knockdown or DN-Mis18 expression. 3xGFP-CENP-A is scaled equivalently. (D) Quantification of 3xGFP-CENP-A fluorescence incorporation relative to control injected oocytes. Points represent the average of all centromeres from individual total oocytes. Error bars represent the mean and standard deviation of at least 29 oocytes. (E) Oocytes following 12 days culture while expressing DN-Mis18 or with control injection, stained for endogenous CENP-C (scaled equivalently), microtubules (scaled non-linearly), and DNA. (F) Graph showing percent of oocytes with normal MI chromosome alignment at the indicated timepoints following a prophase I arrest. Points represent the mean and standard deviation for 3 trials for a total of at least 92 total oocytes per day. Scale bars = 5 μm.

### CENP-A deposition in prophase I arrested oocytes requires the Mis18 complex

We hypothesized that the gradual CENP-A incorporation that occurs in starfish oocytes is required to maintain centromere identity, ensuring a precise first meiotic division after exiting prophase I arrest. To test this premise, we first sought to define the requirements for the deposition of new CENP-A nucleosomes in starfish. Depletion of the Mis18 complex subunit Mis18BP1, a canonical CENP-A deposition factor (McKinley and Cheeseman, 2016), by morpholino injection strongly reduced the G1 deposition of new CENP-A that normally occurs in the female egg pronucleus (Fig. S3C,D). Furthermore, expression of an N-terminally tagged mCherry-Mis18 (hereafter called DN-Mis18) acted in a dominant negative manner to block CENP-A deposition in the egg pronucleus (Fig. S3C,D), likely by preventing Mis18 holocomplex formation. Following fertilization, the resulting Mis18BP1 morphant or DN-Mis18-expressing embryos displayed severe chromosome mis-segregation (Fig S3E), indicative of defective centromere and kinetochore function. These results highlight the important role for the Mis18 complex in maintaining centromere identity in rapidly dividing cells.

We next tested whether the Mis18 complex was required for the gradual CENP-A incorporation that occurs during an extended prophase I arrest. We first depleted arrested oocytes for Mis18BP1 by morpholino or expressed DN-Mis18, and then expressed 3xGFP-CENP-A (Fig. 4C). In contrast to controls, oocytes with Mis18BP1 knockdown or DN-Mis18 expression displayed substantially reduced CENP-A incorporation at centromeres during prophase I arrest (Fig 4C,D). These findings indicate that, in addition to the previously defined role for the Mis18 in cycling cells to promote rapid CENP-A replenishment during G1 (Jansen et al., 2007; McKinley and Cheeseman, 2016), the gradual CENP-A deposition that occurs during the prophase I arrest also requires the canonical CENP-A deposition machinery.

### Prophase I CENP-A deposition is required for faithful meiotic chromosome segregation

The ongoing CENP-A deposition in prophase I-arrested oocytes suggests that active centromere maintenance might be necessary to support the meiotic divisions. To test this hypothesis, we inhibited CENP-A incorporation by expressing DN-Mis18 in prophase I-arrested oocytes. We then assessed meiotic chromosome segregation following hormonal stimulation at different time points. Meiosis proceeded normally at time points following up to 4 days of DN-Mis18 expression, indicating that CENP-A remained at sufficient levels over this arrest period (Fig. 4E,F). However, by 8 days following DN-Mis18 expression, we observed increased chromosome misalignment during meiosis I compared to controls (Fig. 4E,F). The prevalence of chromosome misalignment errors continued to increase with the length of the arrest period (Fig. 4E,F). Importantly, centromeres on misaligned chromosomes in DN-Mis18 embryos contained substantially reduced levels of the CENP-A binding protein CENP-C (Fig. 4E), suggesting a defect in centromere maintenance. Based on the ongoing CENP-A incorporation in prophase I-arrested oocytes (Fig. 4A,B) and the defective meiotic chromosome alignment that occurs when this deposition is prevented (Fig. 4E,F), we conclude that centromeric CENP-A is actively incorporated during the prophase I arrest in oocytes to maintain its full levels at centromeres. In contrast, a CENP-A translation-blocking morpholino failed to reduce endogenous CENP-A levels in prophase I-arrested oocytes after 10 days in culture, despite the fact that this morpholino potently reduced CENP-A levels after these cultured oocytes were then fertilized and allowed to develop for 18 hours (Fig. S3F). These results suggest that CENP-A protein is stable in oocytes (also see Smoak et al., 2016), but that the centromere population is dynamically exchanged with a soluble pool to maintain its full levels at centromeres. These findings present a surprising contrast with previous models in which CENP-A is immobile at centromeres (Jansen et al., 2007).

### Plk1 licensing distinguishes rapid G1 CENP-A deposition from gradual deposition during quiescence

The results described above suggest that there are two distinct, but related CENP-A deposition processes: 1) A rapid pulse of CENP-A deposition that occurs during G1 in cells undergoing mitotic divisions to replenish CENP-A levels following DNA replication, and 2) Gradual CENP-A deposition that occurs in non-dividing cells to maintain CENP-A levels. We next sought to define the basis for this substantial difference in behavior and the rate of CENP-A deposition by determining the requirements that distinguish G1 and G0 deposition. Our work demonstrates that the Mis18 complex is required for both modes of CENP-A deposition. However, our prior work additionally implicated the kinase Plk1 as a licensing factor for CENP-A deposition during G1 required to promote rapid CENP-A deposition (McKinley and Cheeseman, 2014). Surprisingly, in contrast to the clear requirement of Plk1 for the rapid pulse of CENP-A deposition that occurs following mitotic exit (Fig. S4; McKinley and Cheeseman, 2014), we found that treatment with the Plk1 inhibitor BI2536 did not prevent new CENP-A deposition in quiescent RPE-1 cells (Fig. 5A,B). These findings indicate that gradual CENP-A deposition in quiescent cells requires the canonical CENP-A deposition machinery, but not licensing by Plk1, which may promote the more rapid CENP-A deposition observed in G1 cells.

**Figure 5.**
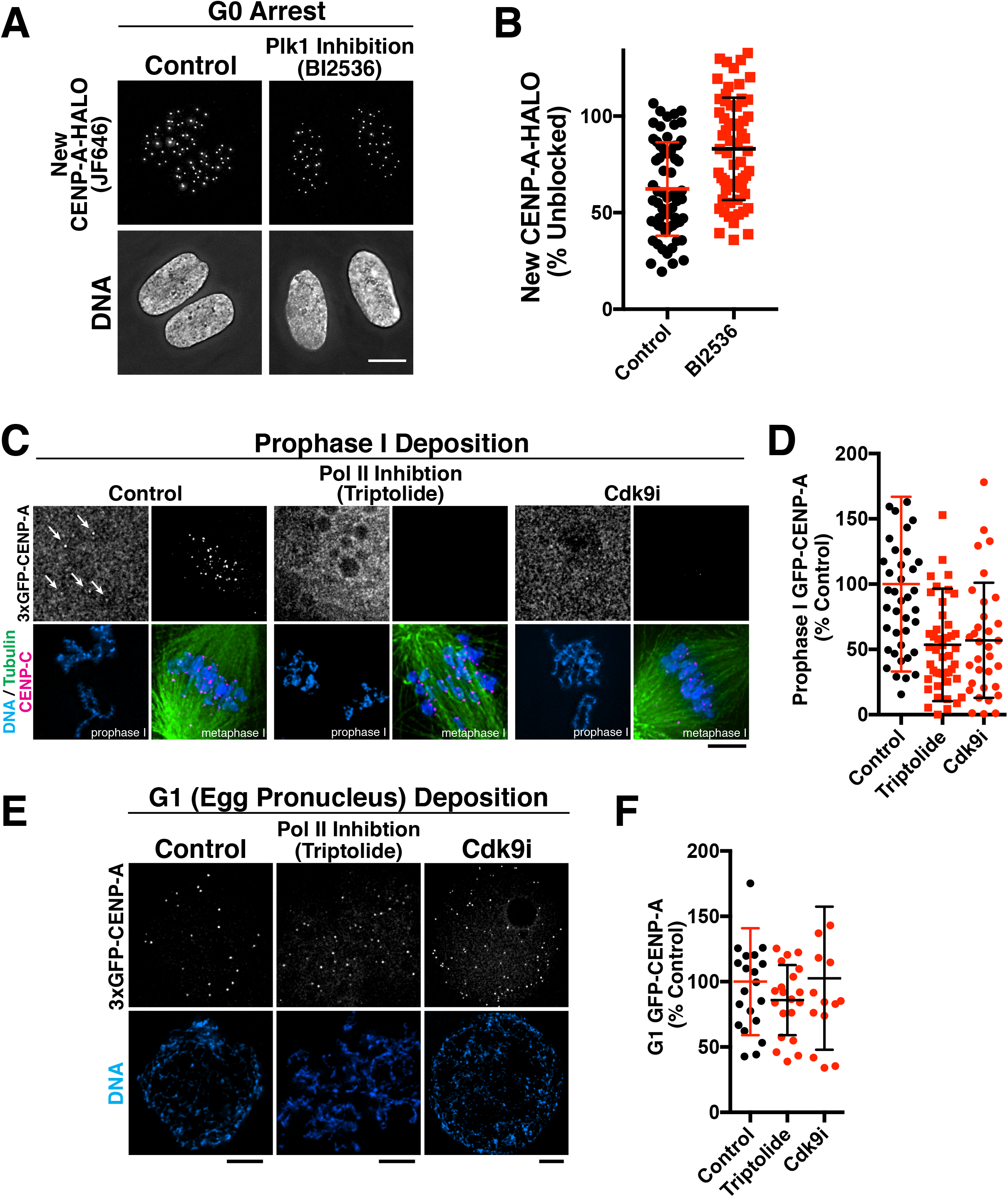
Defining unique requirements for CENP-A deposition during G1 and quiescence. (A) Plk1 is required for G1 CENP-A deposition, but not CENP-A deposition in G0-arrested cells. Immunofluorescence images showing that treatment with 100 nM of the Plk1 inhibitor BI2536 does not prevent incorporation of new CENP-A-Halo in quiescent cells. Scale bar = 5 μm. (B) Quantification of new CENP-A-Halo fluorescent intensity relative to DMSO controls. Some data points from control cells are repeated from Fig. 1C,D. Each point represents the average of all centromeres in a cell. Error bars represent the mean and standard deviation for 62 cells/condition. (C) Transcription promotes CENP-A incorporation in quiescent oocytes. Inhibition of Pol II with 10 μM of triptolide, or Cdk9 with 1 μM of LDC000067 reduces incorporation of new 3xGFP-CENP-A in prophase after 9 days in culture. Scale bar = 5 μm. (D) Quantification of 3xGFP-CENP-A incorporation relative to DMSO control. Each point represents the mean of all centromeres from one oocyte. Error bars represent the mean and standard deviation (control, n = 41; triptolide, n = 47; Cdk9i, n = 33). (E) Inhibition with 10 μM triptolide or 1 μM LDC000067 during prophase I arrest and chronically in meiosis does not alter incorporation of new 3xGFP-CENP-A in G1. Scale bars = 5 μm. (F) Quantification of 3xGFP-CENP-A incorporation relative to DMSO control. Each point represents the mean of all centromeres from one oocyte. Error bars represent the mean and standard deviation (control n = 21, triptolide n = 20, Cdk9i n = 15).

### Blocking transcription reduces new CENP-A deposition in arrested oocytes

The results described above suggest that CENP-A levels are maintained in quiescent cells through balanced loss and incorporation. An important question is what forces evict CENP-A nucleosomes, thereby necessitating their replacement. Growing evidence indicates that centromeres are transcribed by RNA polymerase II (Chan et al., 2012; Grenfell et al., 2016; Saffery et al., 2003), which could promote nucleosome displacement. Therefore, we sought to test whether transcription is required for the ongoing deposition we observed in quiescent cells. Disrupting transcription in G0-arrested human tissue culture cells would have severe consequences for cellular function. In contrast, oocytes have substantial pre-existing mRNA pools such that they do not require ongoing transcription. Indeed, we found that starfish oocytes could be cultured continuously in the presence of the Pol II inhibitor Triptolide or the Cdk9 inhibitor LDC000067 without the loss of viability. The presence of transcription inhibitors did not prevent the translation of 3xGFP-CENP-A following mRNA injection based on cellular fluorescence. However, when transcription was inhibited using these compounds, we found that new CENP-A deposition (based on the incorporation of newly synthesized 3xGFP-CENP-A; see Fig. 4A) was reduced by an average of ~50% in prophase I-arrested oocytes over a 9-day time course (Fig 5C,D). In contrast, when these oocytes incubated in the presence transcription inhibitors for 9 days were allowed to exit meiosis and form the female egg pronucleus, they robustly deposited CENP-A in G1 to levels similar to controls, indicating that the inhibitors did not grossly alter the presence or function of the deposition machinery (Fig 5E,F). These data suggest that Pol II transcription at the centromere provides at least part of the destabilizing force that evicts CENP-A nucleosomes, necessitating their replacement by a Plk1-independent mechanism, but is not generally required for CENP-A deposition.

### Centromere identity is not maintained in terminally differentiated muscle cells

Our work demonstrates that two dramatically different cell types that retain their proliferative potential – oocytes and serum-starved tissue culture cells– display a low rate of CENP-A deposition and exchange. To test whether ongoing CENP-A incorporation is a general feature of non-dividing cells, we next assessed CENP-A levels in terminally differentiated cells. We first tested the mouse C2C12 myoblast-like cell line as an in vitro model for terminal differentiation. C2C12 cells differentiate to form multi-nucleated myotubes upon serum starvation (Yaffe and Saxel, 1977). Following one week in differentiation conditions, a portion of cells remained as mononucleated, quiescent cells. These quiescent reserve cells retained robust centromeric CENP-A and CENP-C (Fig. 6A,B; data not shown). In contrast, the myonuclei in the multi-nucleated myotubes were substantially depleted for both proteins, with fewer centromere foci and less total CENP-A and CENP-C fluorescence (Fig. 6A,B). To determine whether a similar loss of CENP-A occurs in vivo, we conducted immunofluorescence on mouse tissue sections. The mammalian liver has remarkable regenerative capacity and quiescent hepatocytes retain the ability to proliferate into adulthood. We found that adult hepatocyte nuclei from unperturbed liver displayed robust levels of CENP-A (Fig. 6C,D). In contrast, we found that nuclei in two terminally differentiated tissues, including skeletal muscle (gastrocnemius) and heart cardiomyocytes, had significantly reduced levels of CENP-A and fewer detectable centromere foci (Fig. 6C,D). These results suggest that active centromere maintenance is not a default program of all cells, but might occur specifically in cells that need to divide again following quiescence.

**Figure 6.**
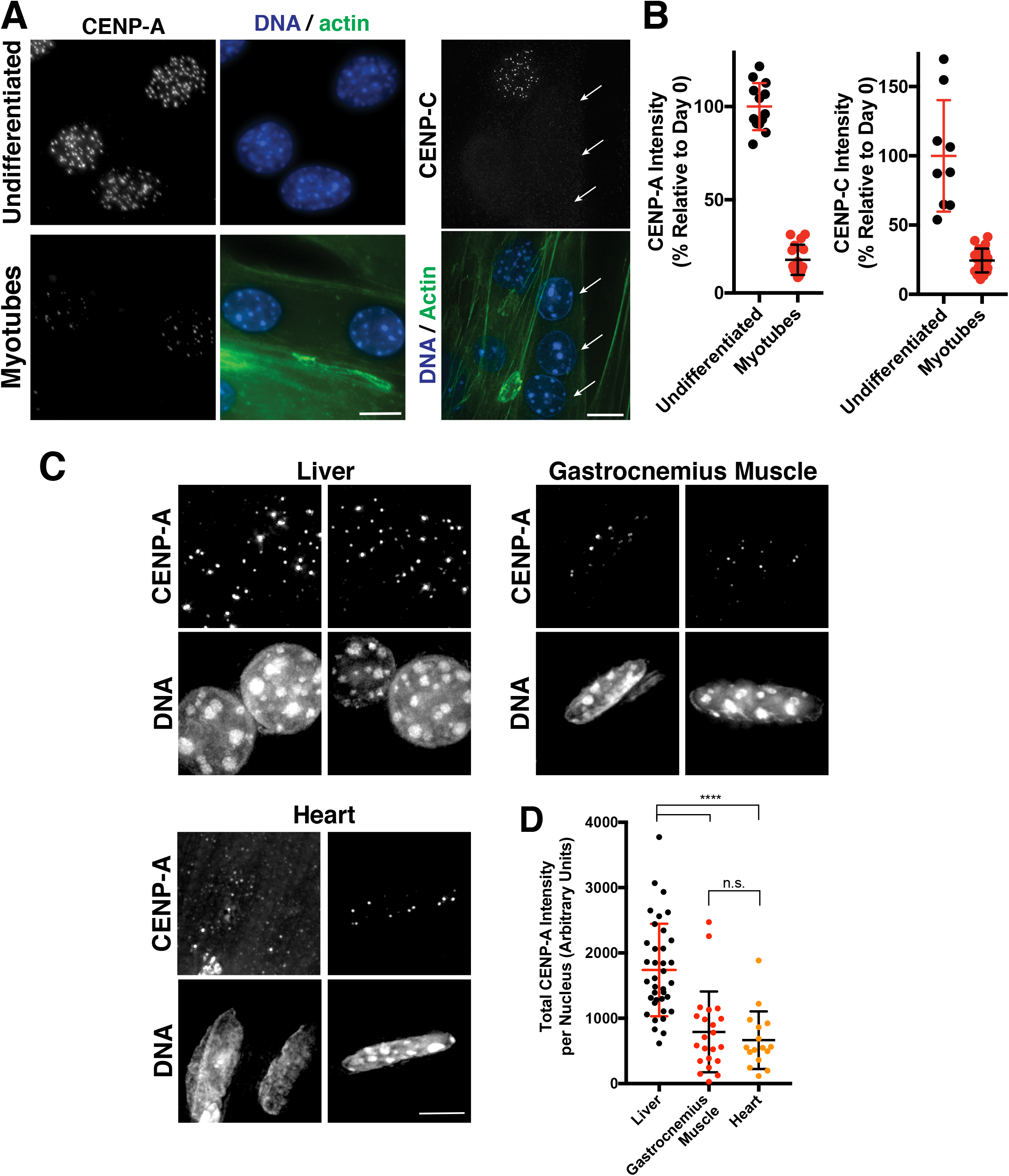
CENP-A levels decline in terminally differentiated muscle cells (A). Immunofluorescence images showing localization of CENP-A (left) and CENP-C (right) in mouse C2C12 cells that are either growing in culture (undifferentiated myoblasts) or were differentiated by serum starvation to form multi-nucleated muscle myotubes. Phalloidin staining is a marker for myotubes. Arrows indicate myotubule nuclei in the CENP-C experiments, whereas the other nuclei with CENP-C staining represents a quiescent, undifferentiated cell. Scale bar = 10 μm. (B) Quantification of CENP-A and CENP-C fluorescent intensity in either undifferentiated C2C12 cells or following myotube formation. (C) Immunofluorescence images of the indicated mouse tissues showing CENP-A levels in vivo. Scale bar = 5 μm. (D) Quantification of CENP-A fluorescent intensity in the indicated tissues (Liver, n=37 nuclei; Gastrocnemius muscle, n = 22; Heart, n=16). For analysis of CENP-A levels in liver hepatocytes, the ploidy of an individual nucleus was determined based on its diameter (see (Knouse et al., 2014)). The intensity of tetraploid nuclei was divided by 2 to compare to the diploid state in muscle cells. **** p < 0.0001

## Discussion

Here, we define a mechanism critical for ensuring the maintenance and inheritance of the epigenetically-defined centromere during extended periods of cellular quiescence. During development, subsets of cells within the body terminally differentiate and permanently exit the cell cycle, whereas slow-cycling or quiescent cells remain dormant, but poised to divide. Arrested cells require a mechanism to maintain the CENP-A epigenetic mark under all circumstances where future proliferation is required. Our results reveal that, in these arrested states, the centromere is actively maintained by a slow rate of ongoing CENP-A deposition. Prior studies in tissue culture cells have focused upon rapidly dividing cells, in which gradual CENP-A incorporation was likely masked by the more rapid G1 deposition. The ability to maintain constant CENP-A levels through balanced loss and incorporation suggests the presence of a homeostatic control mechanism with important implications for genetic inheritance. Although CENP-A-H4 forms a highly stable protein complex whether at centromeres or in a soluble pool (this paper; Smoak et al., 2016), centromere-bound CENP-A is subject to forces such as transcription that destabilize or evict these nucleosomes necessitating its replacement. Thus, CENP-A nucleosomes must be continually incorporated at centromeres to maintain a constant net level. Gradual CENP-A exchange provides a critical means to maintain centromere identity across diverse physiological states. Based on the requirement for an active program to maintain CENP-A at centromeres, we speculate that the presence and levels of CENP-A may provide a novel biomarker for the future proliferative potential of a cell. This gradual exchange is essential to support subsequent chromosome segregation following cell cycle reentry. Therefore, defects in CENP-A deposition in quiescent cells could result in subsequent chromosome segregation abnormalities contributing to tissue dysfunction, infertility/birth defects, or cancer.

### Experimental Procedures

#### Oocyte injection, culture and maturation

*Patiria miniata* were provided by South Coast Bio Marine (http://scbiomarine.com/) and maintained in aquaria at 15°C. Oocytes were extracted as described previously (Wessel et al., 2010). For long-term culture experiments, oocytes were kept in 6 or 12 well plates containing 0.2 micron filtered adult starfish coelomic fluid (CF) supplemented with antibiotics (a stock of 10 mg/ml trimethoprim, 50 mg/ml sulfamethoxazole prepared in DMSO diluted 1:1000 + 100 units/mL penicillin, 100 units/mL streptomycin final). CF was changed every 2–4 days. For transcription studies (e.g. Fig 5), 10 μM triptolide (Pol II inhibitor, Sigma) or 1 μM LDC000067 (Cdk9 inhibitor, Selleck Chemicals) were added to CF and changed every 2 days. For maturation, oocytes were transferred to seawater and 1-methyladenine (Acros Organics) was added to a working concentration of 10 μM. For experiments in which embryos were analyzed, oocytes were fertilized at 60 minutes after 1-methyladenine addition.

#### Construct and antibody generation

Starfish homologs were identified using sequence analysis tools at echionbase.org, and an ovary *de novo* transcriptome described in (Reich et al., 2015). Full-length PmCENP-A and PmCENP-N were cloned as N-terminal 3xGFP fusions in PCS2+ (von Dassow et al., 2009). PmMis18 (DN-Mis18) was cloned as an N-terminal single mCherry fusion in PCS2+8 (Gokirmak et al., 2012). The first 68 amino acids of PmCENP-A, and the first 250 amino acids of PmCENP-C were expressed as a GST-fusions in BL21(DE3) *E. coli*. Protein expression was induced with 0.1 mM IPTG overnight at 18° C. Protein was purified as described previously (McKinley et al., 2015) and antibodies were generated in rabbits (Covance). Antisera were affinity purified against the immunogens using HiTrap NHS-activated columns according to the manufacturer’s instructions (GE Healthcare), then dialyzed into 50% glycerol-PBS.

#### Oocyte knockdown and mRNA expression

For mRNA expression, plasmids were linearized with Noti and transcribed using mMessage mMachine SP6 followed by poly-A tailing (Ambion) and LiCl precipitation. For injections, freshly extracted oocyte-follicle cell complexes were injected horizontally in Kiehart chambers with 2–4 picoliters of mRNA or morpholino solution in water. 3xGFP-CENPA mRNA was injected at 50–85 ng/μl, whereas 3xGFP-CENP-N and DN-Mis18 was injected at 500 ng/μl in water. Oocytes were then incubated overnight in filtered sea water for rapid assessment of localization, or for up to two weeks in CF for long term perturbations. Custom translation-blocking morpholino antisense oligos were synthesized to CENP-A (5’-TCCGAGCCATGTTCCAAAACAACCT-3’) and MIS18BP1 (5’-GTTGGTTGTGAGAATGTCCCTCCAT-3’) by Gene-Tools. Morpholinos were injected at 500 μM in water. Oocytes were fixed for 3 hours to overnight in a buffer modified from (von Dassow et al., 2009) containing 2% paraformaldehyde, 0.1% Triton X-100, 100 mM Hepes, pH 7.0, 50 mM EGTA, 10 mM MgSO4, and 400 mM dextrose. Oocytes were blocked for 15 minutes in AbDil (3% BSA, 1 X TBS, 0.1% triton X-100, 0.1% Na Azide). Oocytes were stained overnight at 4° C with the anti-CENP-C antibody at 1μg/ml. Microtubules were stained with 1:1000 DM1α (Sigma). GFP booster was used 1:500 to amplify the signal from GFP expression (Chromotek). DNA was visualized using Hoechst. Oocytes were imaged using a DeltaVision Core microscope (Applied Precision/GE Healthsciences) with a CoolSnap HQ2 CCD camera and a 100x 1.40 NA Olympus U-PlanApo objective. Images were deconvolved and maximally projected. For image quantification, all images were acquired using the same microscope and detector settings. Centromere intensities for 3xGFP-CENP-A and 3xGFP-CENP-N were measured using Fiji (Schindelin et al., 2012). Individual centromeres were selected with 5 pixel diameter circles and the total integrated intensity was measured. Background correction was performed by selecting a nearby non-centromeric region of equal size for each centromere and subtracting its integrated intensity from that of the centromere region. The average of all centromeres was then determined per oocyte.

#### Cell line generation and culture

Inducible knockouts for Mis18β, CENP-A, and HJURP were created as described previously (McKinley and Cheeseman, 2017). Briefly, sgRNAs were stably expressed by lentiviral infection in RPE-1 cells harboring inducibly expressible Cas9. Cells were then selected with 10 μg/ml puromycin for 2 weeks. Knockout was induced by addition of 1 μg/ml doxycycline for two days. For endogenous tagging of CENP-A with the Halo tag (cKC288), RPE-1 cells were co-transfected with pX330 containing CAS9 and an sgRNA targeting 3’ UTR of the genomic CENP-A locus (TGTCATCAGTGTTGCCATCG) and a repair CENP-A-HALO fusion-template pKC158, which harbors the HaloTag and a Neomycin resistance cassette derived from pL452 flanked by approximately 1 kb homology arms (5’-arm primers AGTCGGCGGAGACAAGGGTGGCTAAA, GCCGAGTCCCTCCTCAAGGCCCCGGATCCTCCGGGCC, 3’-arm primers GCTCCTGCACCCAGTGTTTCTGTCA, ACATCCGTTGACAAGCACAGTC) resistant to CAS9 by mutation of the PAM sequence TGG to TAA. Cells were then selected with 1 mg/mL G418 (Gibco) before clonal cell line generation. The resulting cell line was homozygously tagged as determined by western blot for CENP-A.

Cells were cultured in Dulbecco’s modified Eagle medium (DMEM) with 10% tetracycline-free fetal bovine serum, 100 units/mL penicillin, 100 units/mL streptomycin, and 2 mM L-glutamine (Complete Media) at 37°C with 5% CO_2_. For induction of quiescence, RPE-1-derived cell lines were allowed to reach confluence, and then shifted to Complete Media as above but with 0.1% fetal bovine serum. To ensure that no mitotic cells were present in the population, the media was supplemented with 10 μM STLC where indicated. For cells in quiescence, the media containing 0.1% serum was changed daily.

C2C12 myoblasts were cultured as previously described (Lawson and Purslow, 2000). In brief, proliferating myoblasts were grown in in DMEM with 10% FBS and 1% penicillin/streptomycin. For immunofluorescence, myoblasts were seeded onto uncoated glass coverslips and grown to confluence. To induce differentiation, the myoblasts were shifted to differentiation medium (DMEM with 10% horse serum). Myotubes were fixed in methanol at 20° C for 20 minutes. Coverslips were washed three times in PBS, and then blocked and permeabilized in PBS containing 0.1% Triton X-100 and 2% BSA. Primary antibodies to mouse CENP-A (Cell Signaling Technology, C51A7 Rabbit mAb #2048; diluted 1:1000) or mouse CENP-C (Cheeseman lab) in block solution and incubated for 1 hour at room temperature. The coverslips were then washed three times in PBS and then incubated for 1 hour with 1:250 Alexa567 conjugated anti rabbit-secondary antibodies. The cells were then counterstained with DAPI and phalloidin 688.

#### HaloTag pulse-chase detection and antibody staining in RPE-1 cells

To assess CENP-A dynamics in quiescence, cKC288 cells were seeded onto uncoated coverslips and grown to confluence. The cells were then shifted to 0.1% serum-containing media and cultured for 5 days to ensure complete mitotic exit. The cells were then either left unblocked or blocked with 125 nM JF549-HaloTag ligand (gift from Luke Lavis; (Grimm et al., 2017; Grimm et al., 2015)). Coverslips were then taken at periodic intervals and fixed with 100% ice-cold methanol. New CENP-A was visualized by staining coverslips with 30 nM JF646-HaloTag ligand (gift from Luke Lavis; (Grimm et al., 2017)). For immunofluorescence, coverslips were washed with PBS containing 0.1% Triton X-100, then blocked with AbDil. Anti CENP-A antibody (Abcam, ab13939) was used at 1:1000, and anti-centromere antibodies (ACA) were used at 1:100 (Antibodies, Inc.). Cells were imaged using a DeltaVision Core microscope (Applied Precision/GE Healthsciences) with a CoolSnap HQ2 CCD camera and a 60x 1.40 NA Olympus U-PlanApo objective. Centromere pixel intensities were quantified using Fiji (Schindelin et al., 2012). Z-stacks were projected as single two-dimensional images and centromeres were identified using either the ACA or anti CENP-A antibody signal. The raw integrated density of fluorescence at each centromere within a nucleus was collected and summed as one data point. At least 25 nuclei were analyzed for each time point and condition.

To calculate the percent new CENP-A-HALO, the average JF646 fluorescence per nuclei in unblocked cells was used as total CENP-A. To account for natural differences in CENP-A levels between cells, the anti CENP-A antibody fluorescence was measured for both blocked and unblocked cells at the same location as JF646 signal. The final equation used was as follows:

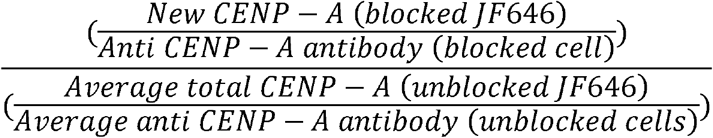

#### Plk1 inhibition studies

To test the requirement of Plk1 for CENP-A deposition, cells were synchronized by inducing quiescence as described above by 3 days culture in 0.1% FBS. Pre-existing CENP-A was blocked using 125 nM JF549-HaloTag ligand, then re-plated onto uncoated coverslips and returned to growth conditions in 10% FBS. After 24 hours (sufficient time to reenter the cell cycle), the cells were arrested in mitosis with 10 μM STLC overnight. The cells were released into complete media without STLC and, after 1.5 hours, treated with either DMSO or 100 nM BI 2536. The cells were then fixed in methanol and chased with 30 nM JF646-HaloTag ligand. In quiescence, 100 nM BI 2536 was added to 0.1% serum-containing media and changed daily. Coverslips were collected and analyzed as above.

#### Mouse Tissue Immunostaining

C57BL/6J mice were purchased from the Jackson Laboratory. All mice were group-housed with a 12-hour light-dark cycle (light from 7 AM to 7 PM, dark from 7 PM to 7 AM) in a specific-pathogen-free animal facility with unlimited access to food and water. Liver, gastrocnemius, and heart were harvested from 8-week-old adult male mice. All animal procedures were approved by the Massachusetts Institute of Technology Committee on Animal Care. Tissues were fixed in 4% paraformaldehyde in PBS at room temperature for 16–24 hours. Fixed tissues were washed with PBS, cryoprotected with 30% sucrose, and frozen in O. C. T. Compound (Tissue-Tek). Slides with 15 μm-thick sections were prepared using a cryostat. Slides were dried at room temperature for 4–24 hours. Slides were rehydrated in PBS for 5 minutes and incubated in sodium citrate buffer (10 mM tri-sodium citrate dihydrate, 0.05% Tween 20, pH 6.0) in a pressure cooker (Instant Pot) for 20 minutes at high pressure. Slides were washed with PBS, dried briefly, and sections outlined with a hydrophobic pen. Sections were incubated with extraction buffer (1% Triton X-100 in PBS) for 15 minutes followed by incubation in blocking solution (3% BSA, 0.3% Triton X-100 in PBS) for 1 hour at room temperature. Sections were incubated with primary antibodies (anti-mouse CENP-A; C51A7 from Cell Signaling) diluted in blocking solution at room temperature for 16–24 hours. Sections were washed three times with blocking solution for 10 minutes each and then incubated with secondary antibodies diluted in blocking solution with 5 μg/mL Hoechst 33342 (Thermo Fisher Scientific) at room temperature for 1–2 hours. Sections were washed with blocking solution for 5 minutes twice and once with PBS for 5 minutes. Sections were mounted with ProLong Gold Antifade Reagent (Life Technologies).

## Acknowledgements

We thank the members of the Cheeseman laboratory for support, input, and critical reading of the manuscript, Terry Orr-Weaver, Peter Reddien, Priya Budde, and Kara McKinley for critical reading of the manuscript, and Luke D. Lavis for JF-HALO-reagents. This work was supported by grants from The Harold G & Leila Y. Mathers Charitable Foundation and the NIH/National Institute of General Medical Sciences (GM088313 and GM108718) to IMC, and an American Cancer Society post-doctoral fellowship to SZS.

## Author Contributions

Conceptualization – SZS, IMC; Methodology – SZS, LSM, KCS, KAK; Validation – SZS, LSM, KCS; Investigation – SZS, LSM, KCS, AP; Writing – Original Draft Preparation – SZS, IMC; Writing – Review & Editing – SZS, LSM, KCS, IMC, KAK; Visualization – SZS, LSM, KCS, IMC; Supervision – PSM, IMC; Funding Acquisition: IMC.

## Supplemental Figure Legends

**Figure S1.**
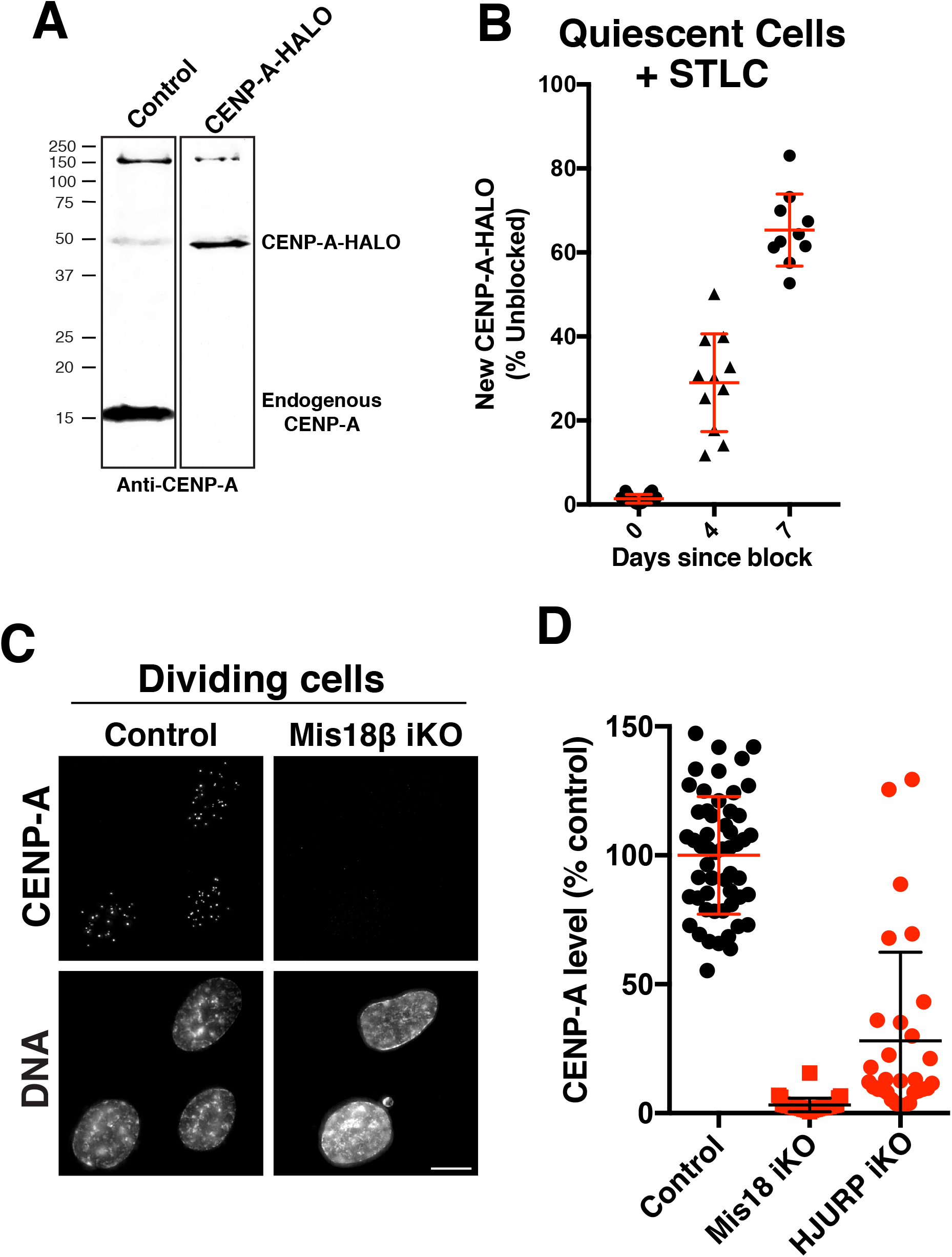
Analysis of CENP-A incorporation. Related to Figure 1 and 2. (A) Western blot showing CENP-A in either control RPE-1, or RPE-1 cells with a HaloTag introduced at the CENP-A C-terminus at the endogenous locus using CRISPR/Cas9. The absence of a band for endogenous CENP-A indicates that both CENP-A loci are tagged. (B) Halo-tag CENP-A incorporation assay in quiescent cells as in Fig. 3C,D, but in cells treated continuously with the Kif11 inhibitor STLC to block any rare mitotic divisions. Graph shows quantification of new CENP-A-Halo fluorescent intensity relative to pre-existing CENP-A. Points represent the sum of all centromeres of individual cells. Error bars represent the mean and standard deviation of at least 10 cells per condition/timepoint. (C) Immunofluorescence images showing CENP-A levels in dividing (non-quiescent) cells either in control RPE-1 or a Mis18β Cas9-based inducible knockout 7 days following Cas9 induction. (D) Quantification of CENP-A intensity relative to control cells from (C) (control, n=54; Mis18 β iKO, n=34; HJURP iKO, n=30).

**Figure S2.**
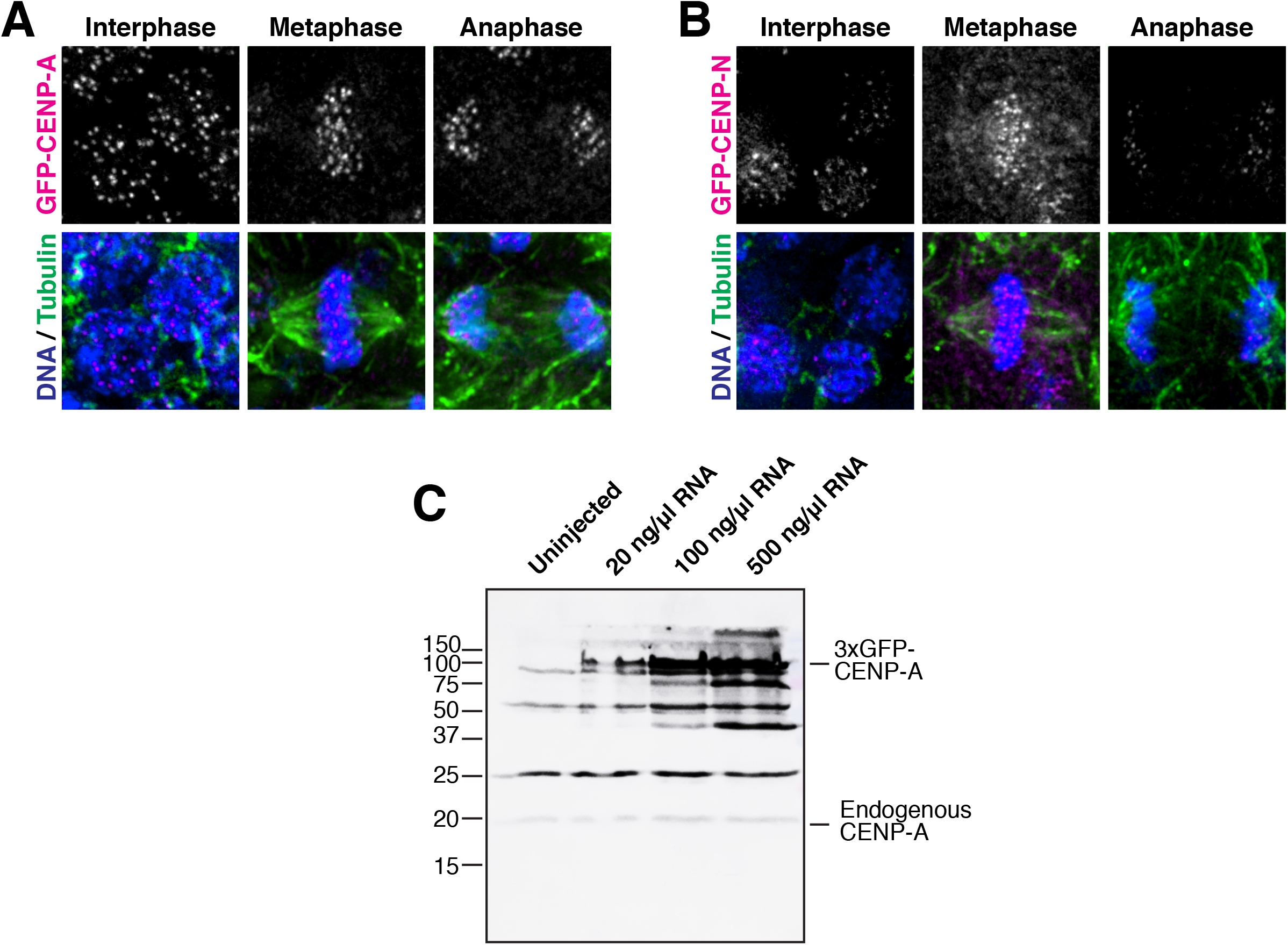
Localization of GFP fusions in oocytes and embryos. Related to Figure 3. (A,B) 3xGFP-CENP-A and CENP-N localize to centromeres in dividing ectodermal cells. Constructs were introduced into prophase I-arrested oocytes by mRNA injection and visualized following fertilization in 18-hour old embryos. CENP-A and CENP-N images are scaled equivalently across cell cycle stages. Scale bars = 5 μm. (C) Western blot for CENP-A in arrested oocytes 18 hours after injection of the indicated concentrations of mRNA or uninjected controls.

**Figure S3.**
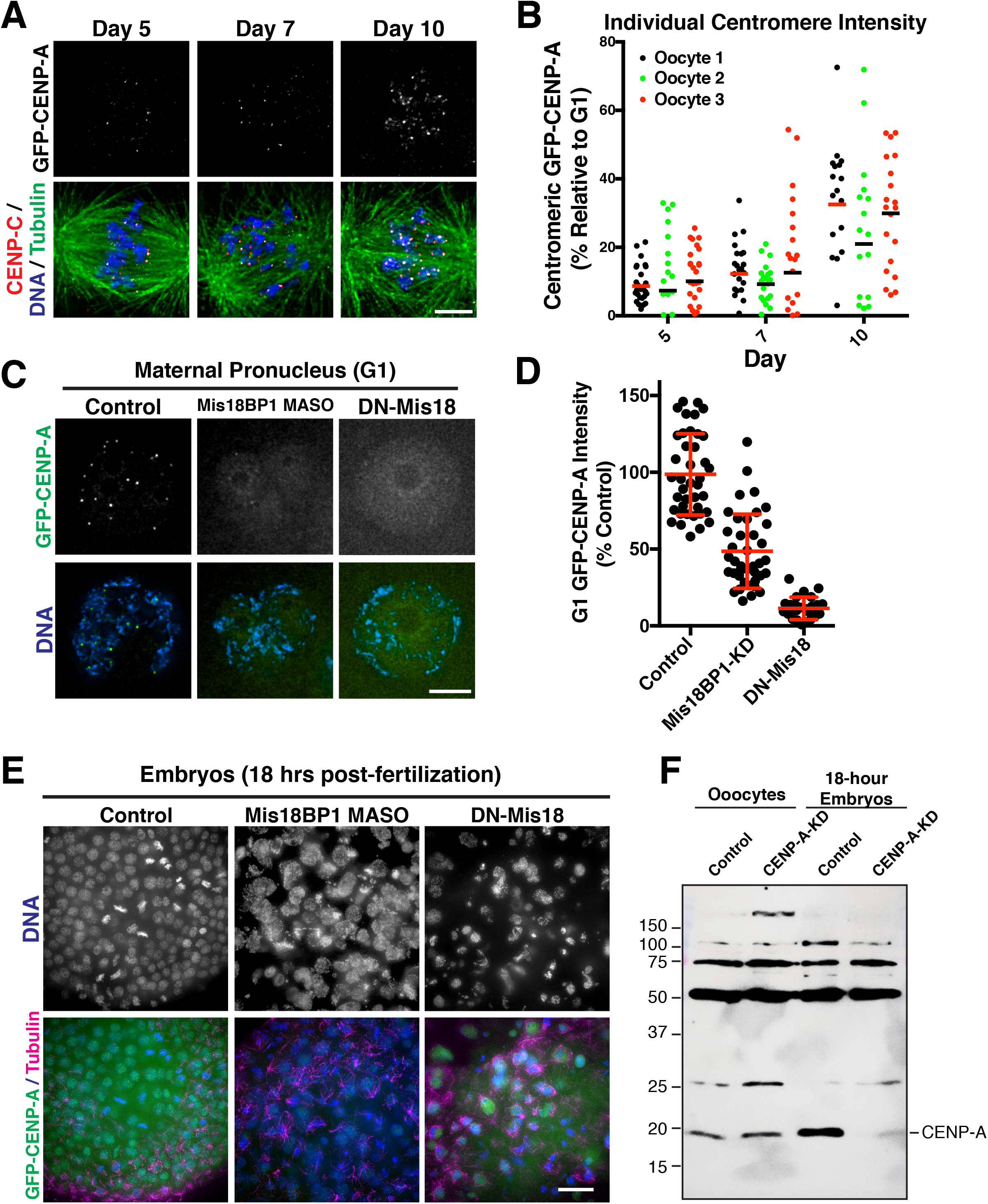
CENP-A protein is stable but gradually deposited in arrested oocytes. Related to Figure 4. (A) CENP-A displays stochastic incorporation at some centromeres over time. Immunofluorescence images showing prophase I deposited CENP-A visualized after meiotic entry for clarity. Scale bar = 5 μm. (B) 3x-CENP-A levels at individual centromeres (each point represents one centromere) from three representative oocytes from the average centromere data presented in Fig, 2b. (C) G1 replenishment of 3x-GFP-CENP-A is reduced following Mis18BP1 knockdown or DN-Mis18 expression in starfish oocytes. Prophase arrested oocytes were injected with mRNA encoding 3xGFP-CENP-A either alone, or together with a Mis18BP1 morpholino (MASO) or dominant negative Mis18 construct (DN-Mis18) Scale bar = 5 μm. (D) Quantification of 3xGFP-CENP-A levels at centromeres in the G1 female pronucleus (from c). Each point represents the average of all identifiable centromeres in one egg. Error bars represent the mean and standard deviation (control, 40 eggs; Mis18BP1 knockdown, 39 eggs; DN-Mis18, 26 eggs). (E) CENP-A incorporation and chromosome segregation are disrupted in 18-hour embryos following Mis18BP1 knockdown or DN-Mis18 expression. Scale bar = 20 μm. (F) Endogenous CENP-A protein is stable in prophase I-arrested oocytes. Anti-CENP-A Western blot for oocytes 10 days after injection of control or CENP-A translation blocking morpholino to test CENP-A levels in prophase I-arrested oocytes, or following 18 hours of embryogenesis.

**Figure S4.**
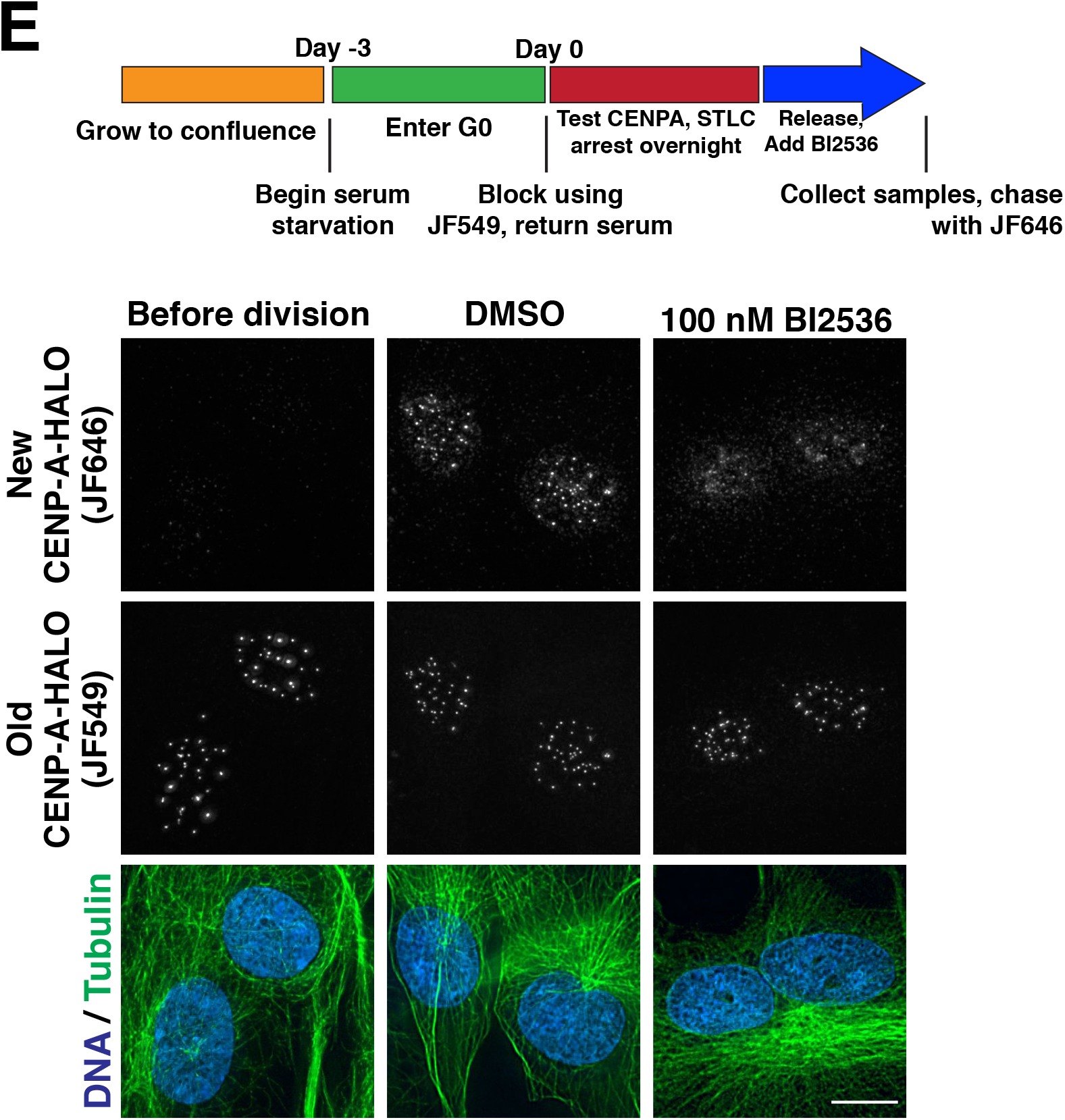
Plk1 is required for G1 CENP-A deposition. Related to Figure 5. (A) Plk1 inhibition blocks new CENP-A deposition during G1. Top, schematic showing growth and Halo block conditions to test new CENP-A deposition in CENP-A-Halo Rpe1 cells. Bottom, immunofluorescence images showing new vs. old CENP-A staining either before division (18 hours following serum addition) or during G1 in either a DMSO control or in the presence of 100 nM of the Plk1 inhibitor BI2526. Scale bars = 10 μm.

